## Supplementary material for "Host diet and evolutionary history explain different aspects of gut microbiome diversity among vertebrate clades"

### Supplemental Tables

All supplemental tables are provided as separate Excel files

**Supplementary Table 1.** Metadata for all samples in the dataset.

**Supplementary Table 2.** Pearson correlation coefficients comparing LIPA values between OTUs with significant local phylogenetic signal (See Fig. 4).

### Supplemental Figures

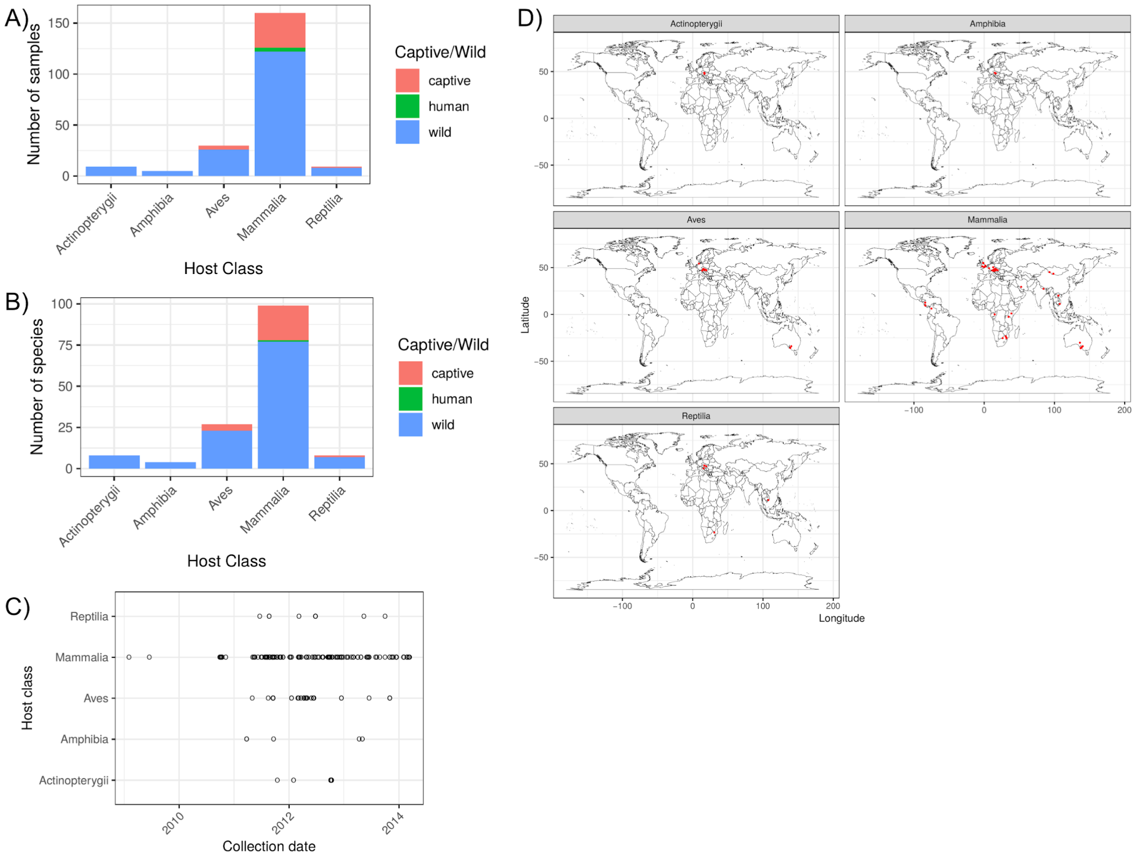


**Supplementary Fig. 1**. *The dataset includes gut microbiome samples spanning 5 host classes, ~5 years, and 6 continents.* A) the number of samples per host class B) the number of species per host class C) the time point of sample collection D) the geographic location of each sample collection.

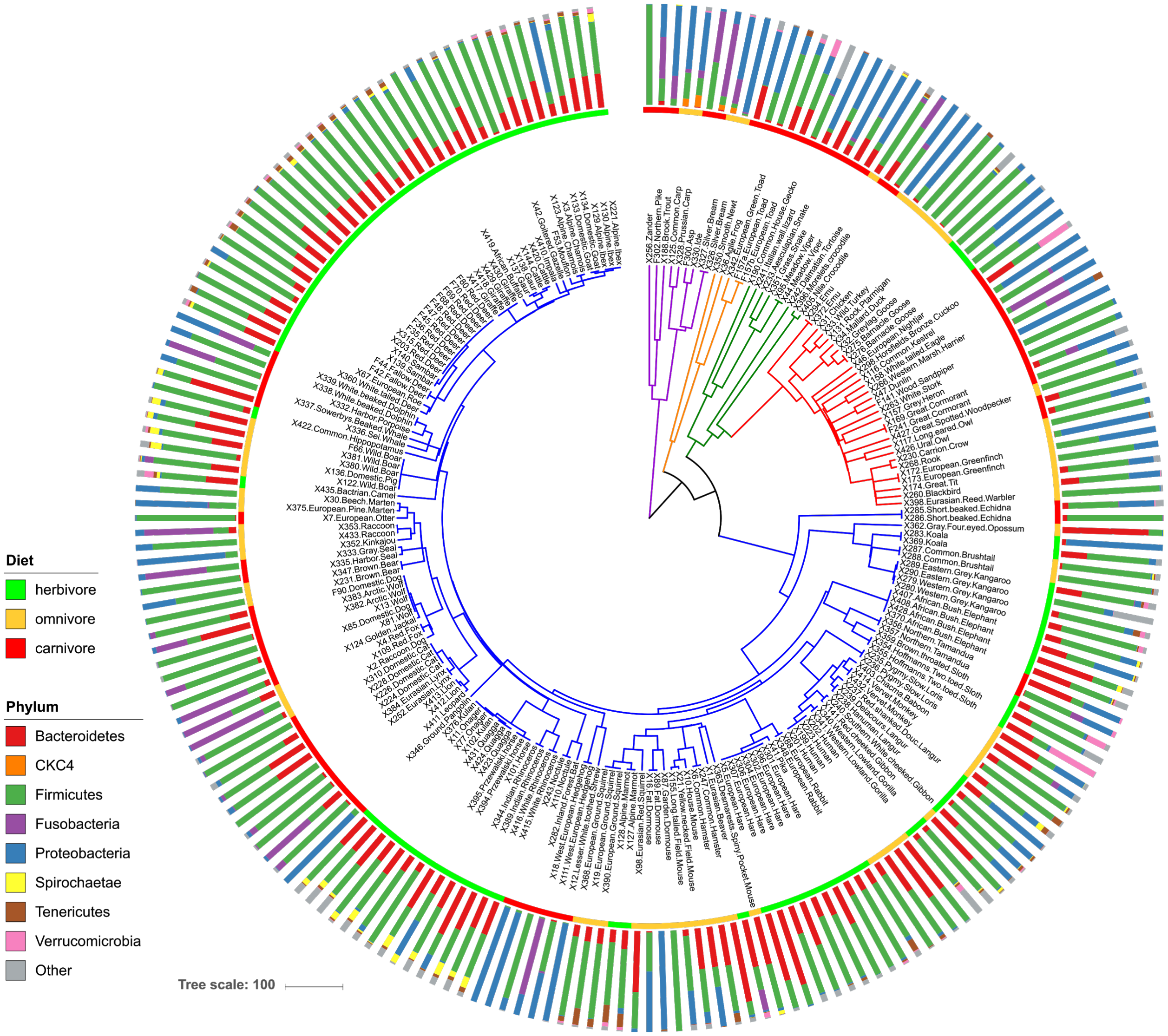


**Supplementary Fig. 2.** *Phylum-level grouping of microbiome diversity by host phylogeny and host metadata*. The host phylogeny is that same as shown in Fig. 1, except tips have been expanded to include all individuals of each species. Host diet and relative abundances of microbial phyla are mapped onto the tree.

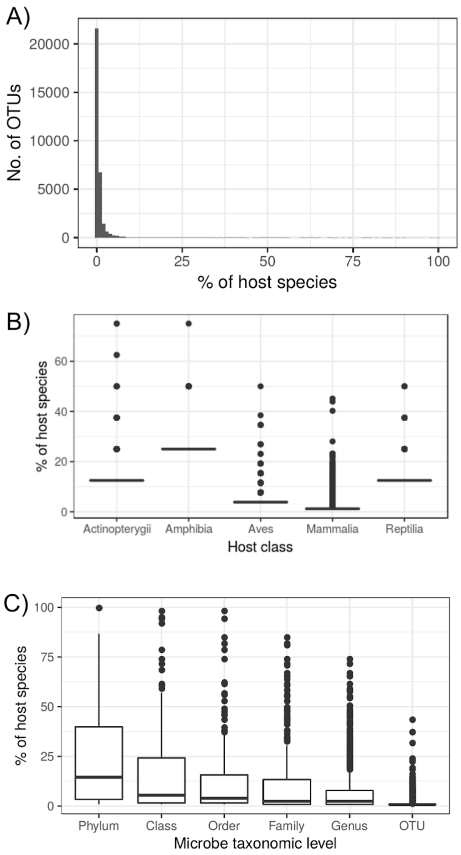


D)

| Taxonomic level | Group | Prevalence |
|---|---|---|
| Phylum | Firmicutes | 100 |
| Phylum | Proteobacteria | 100 |
| Class | Firmicutes;Clostridia | 98.4 |
| Order | Firmicutes;Clostridia;Clostridiales | 98.4 |
| Class | Proteobacteria;Betaproteobacteria | 95.3 |
| Class | Proteobacteria;Gammaproteobacteria | 94.5 |
| Order | Proteobacteria;Betaproteobacteria;Burkholderiales | 94.5 |
| Class | Firmicutes;Bacilli | 92.2 |
| Phylum | Actinobacteria | 86.7 |
| Phylum | Bacteroidetes | 85.9 |
| Order | Proteobacteria;Gammaproteobacteria;Enterobacteriales | 85.2 |
| Family | Proteobacteria;Gammaproteobacteria;Enterobacteriales; Enterobacteriaceae | 85.2 |
| Order | Firmicutes;Bacilli;Lactobacillales | 82 |
| Family | Firmicutes;Clostridia;Clostridiales;Lachnospiraceae | 82 |
| Family | Firmicutes;Clostridia;Clostridiales;Clostridiaceae 1 | 81.2 |
| Class | Bacteroidetes;Bacteroidia | 78.9 |
| Order | Bacteroidetes;Bacteroidia;Bacteroidales | 78.9 |

**Supplementary Fig. 3**. *OTUs are sparsely distributed in the dataset*. A) OTU prevalence across all host species (found in at least one individual). B) OTU prevalence across host species, grouped by host class. C) Microbe taxonomic group prevalence across all host species. D) Microbe taxonomic groups with a prevalence of >75 % across all host species.

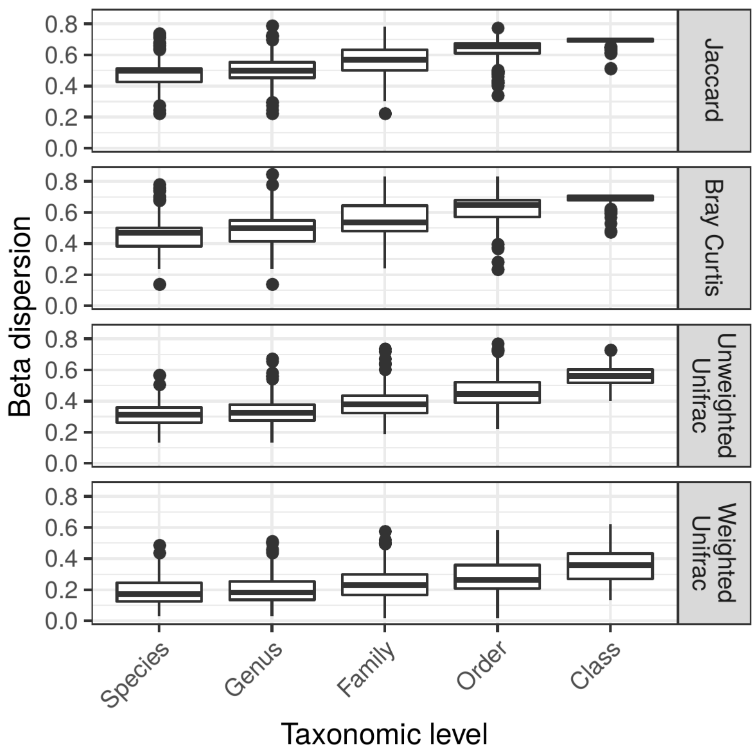


**Supplementary Fig. 4.** *Beta-diversity is more constrained at finer taxonomic levels.* The boxplots show the distribution of beta-dispersion (distance from multivariate centroids of the group), with groups defined as host clades at each taxonomic level.

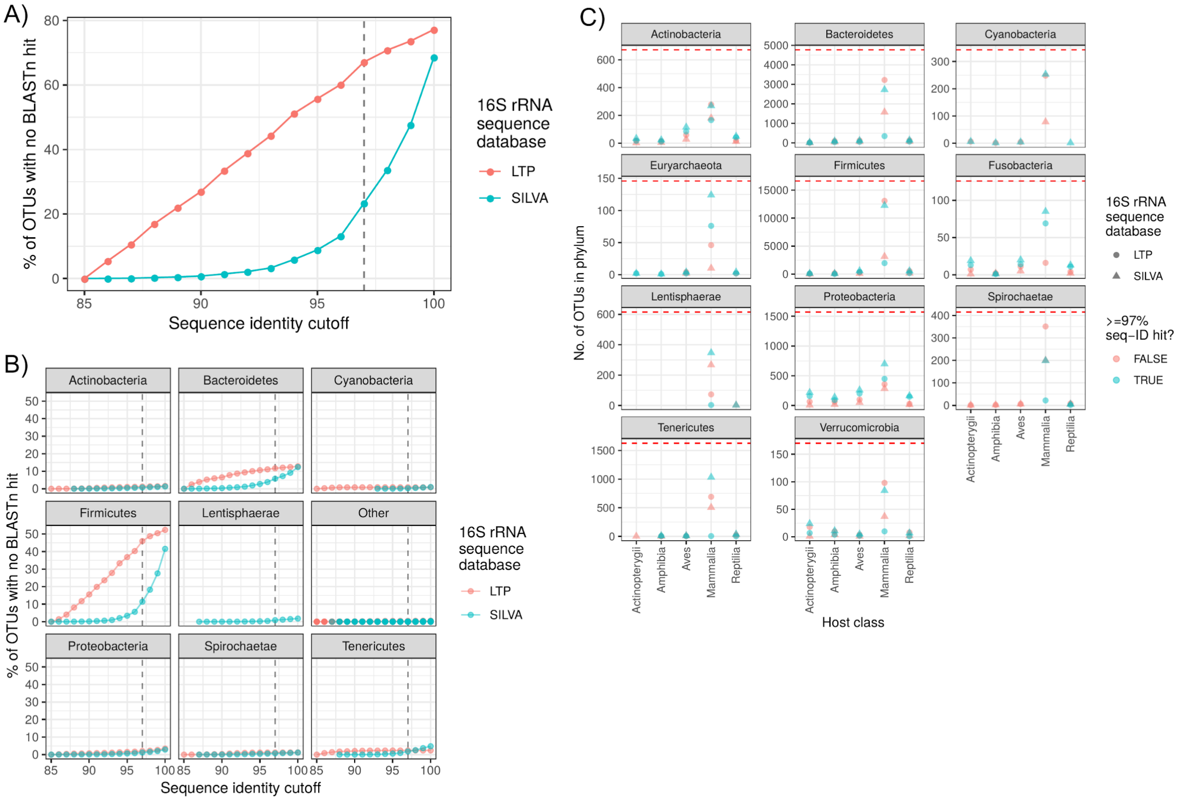


**Supplementary Fig. 5.** *Many of the OTUs in the dataset lack (un)cultured representatives.* The plots show the distribution of BLASTn hits between OTU representative sequences and 16S rRNA sequences of cultured representative taxa in the SILVA All Species Living Tree database ("LTP") or the entire SILVA database de-replicated at 99 % sequence identity ("SILVA"). The vertical dashed line in A) and B) signifies a percent sequence identity of 97 %. "Other" in B) comprises all other phyla. ">=97% seq-ID hit?" in C) signifies whether the OTU has at least one BLASTn hit to a taxon in either SILVA database with ≥97 % sequence identity. For each category in C), only OTUs with a prevalence of >0 are counted, while the dashed line signifies all OTUs in that phylum. Only phyla with >100 OTUs are shown in C).

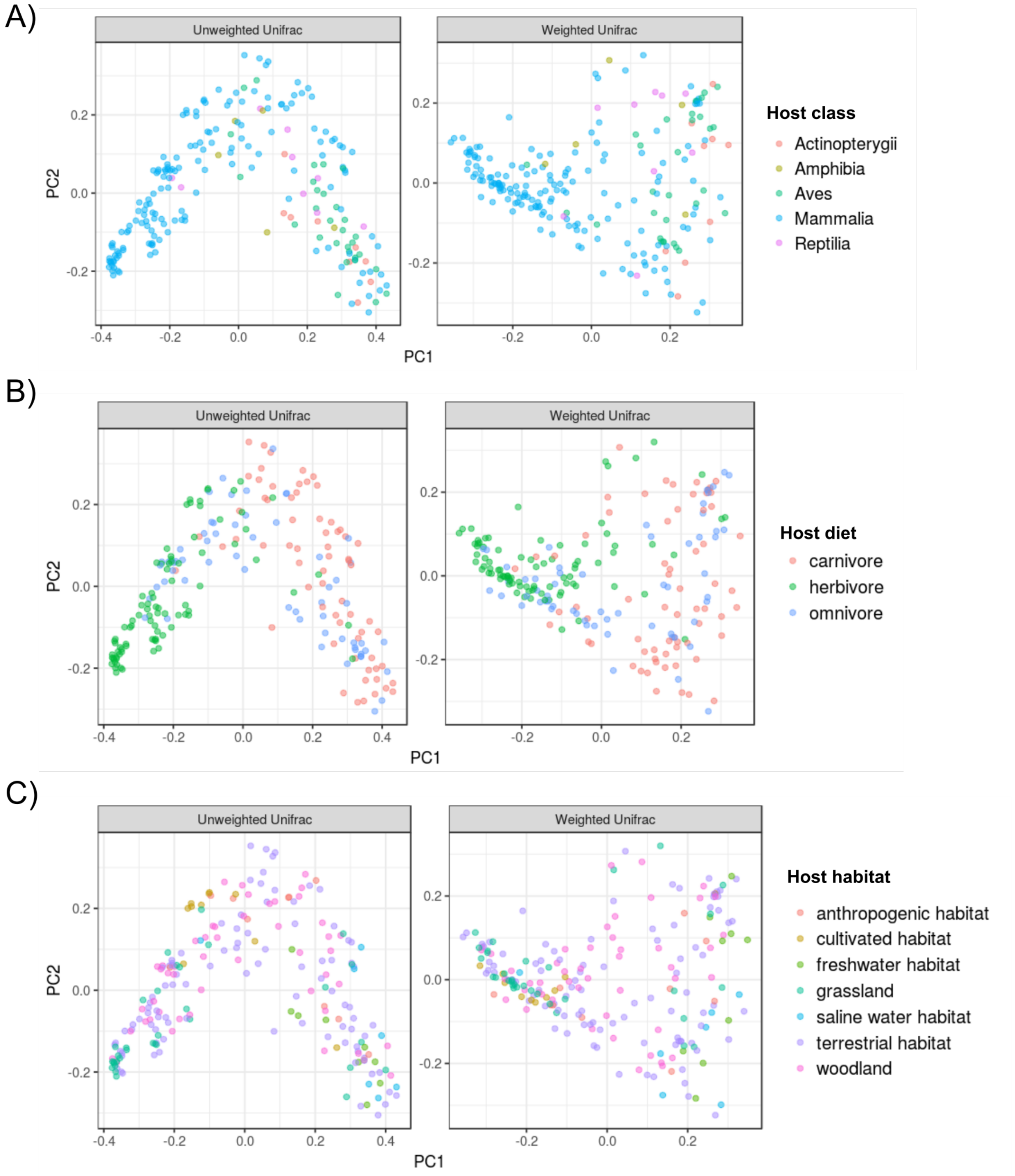


**Supplementary Fig. 6**. *PCoA plots* of *microbiome beta-diversity reveal some grouping by host taxonomy and diet*. Principal coordinates (PCoA) ordinations of unweighted and weighted Unifrac distances among all samples, with samples colored by host A) class B) diet C) habitat. The variance explained by PC1 and PC2 of the unweighted Unifrac PCoA is 17 and 6 %, respectively. The variance explained by PC1 and PC2 of the weighted Unifrac PCoA is 26 and 11 %, respectively.

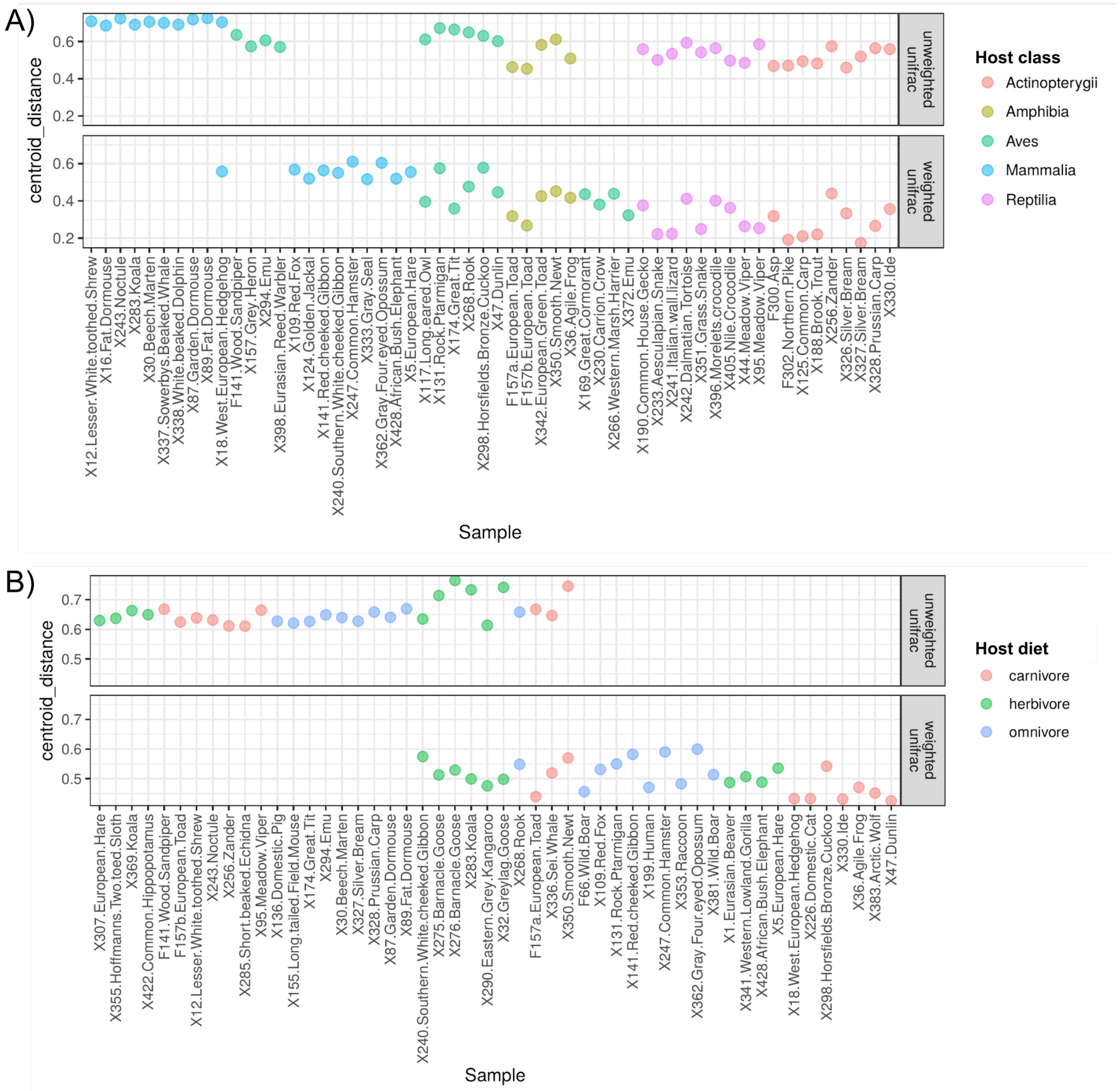


**Supplementary Fig. 7.** *Outlier samples differ among beta-diversity metrics and groupings*. Centroid distance of samples for unweighted or weighted Unifrac, with centroids defined by host taxonomy (A) or host diet (B). Only samples with a centroid distance of >0.4 (ie., the largest outliers) are shown.

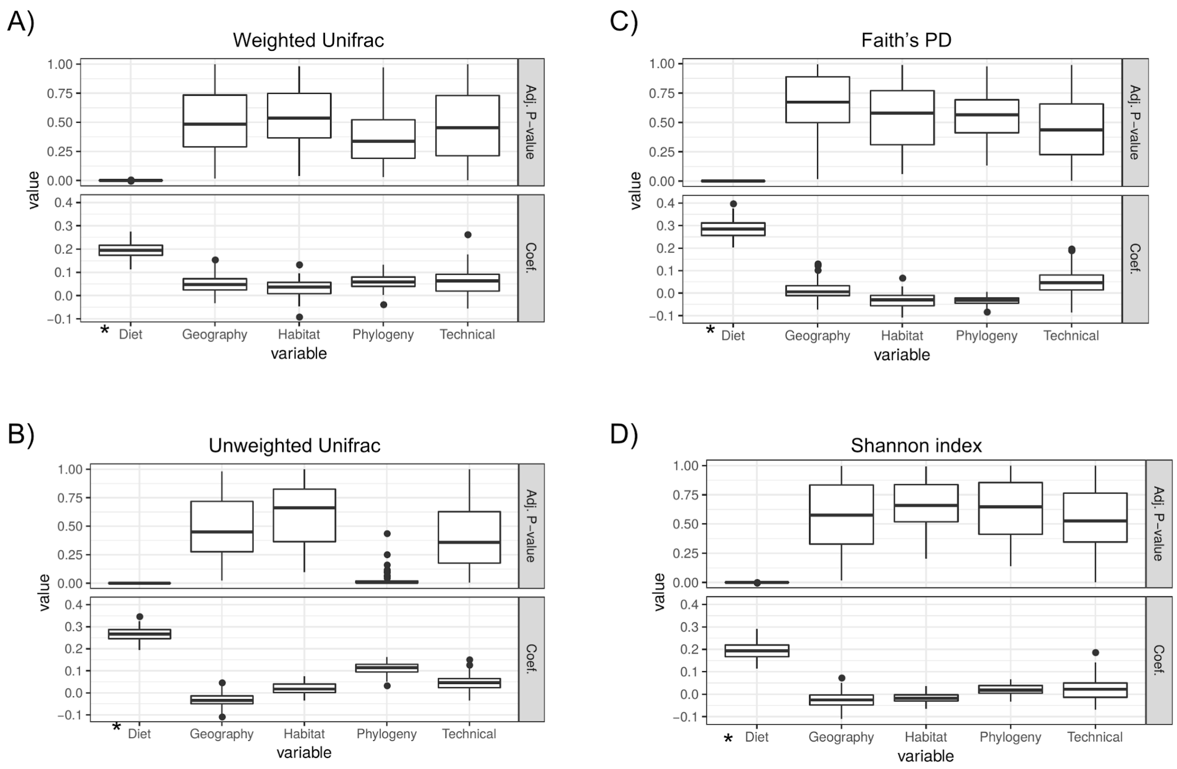


**Supplementary Fig. 8.** *Very similar MRM results obtained if just selecting just one sample per family, which reduces sample size biases towards Mammalia*. The figure is the same as Fig. 2, but for each dataset subset, only one sample was selected per family instead of per species. "*" signifies significance (Adj. p-value <0.05 for ≥95 % of dataset subsets; see Methods).

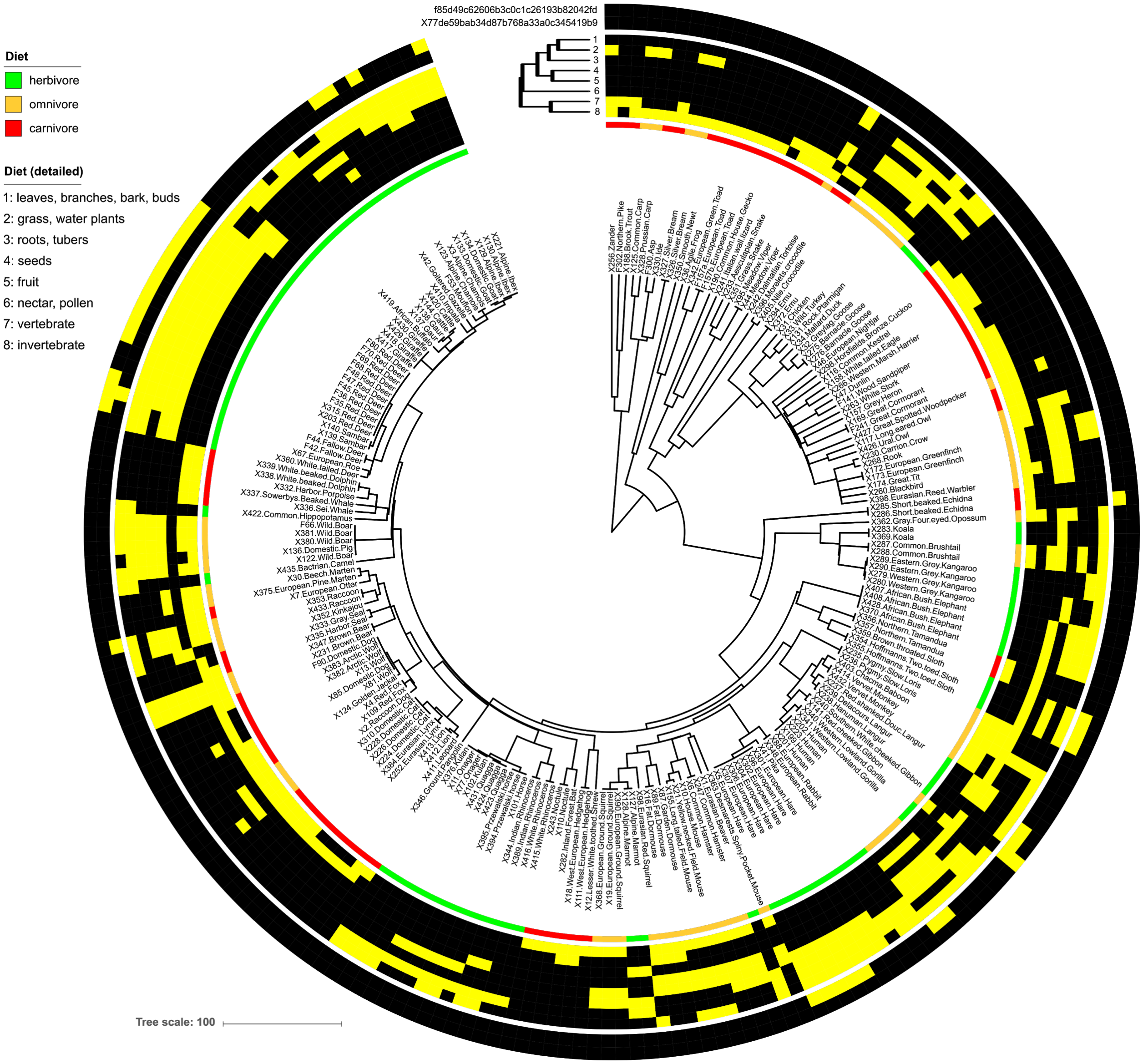


**Supplementary Fig. 9.** *OTUs significantly explained by diet differ in their prevalence among hosts and diets.* The phylogeny is the same as shown in Supplementary Fig. 2. The presence of the PGLS-significant OTUs (See Fig. 3) are mapped onto the host tree (outer ring) along with the detailed host diet characteristics (middle ring) and the general diet (inner ring). Yellow and black squares signify presence and absence, respectively.

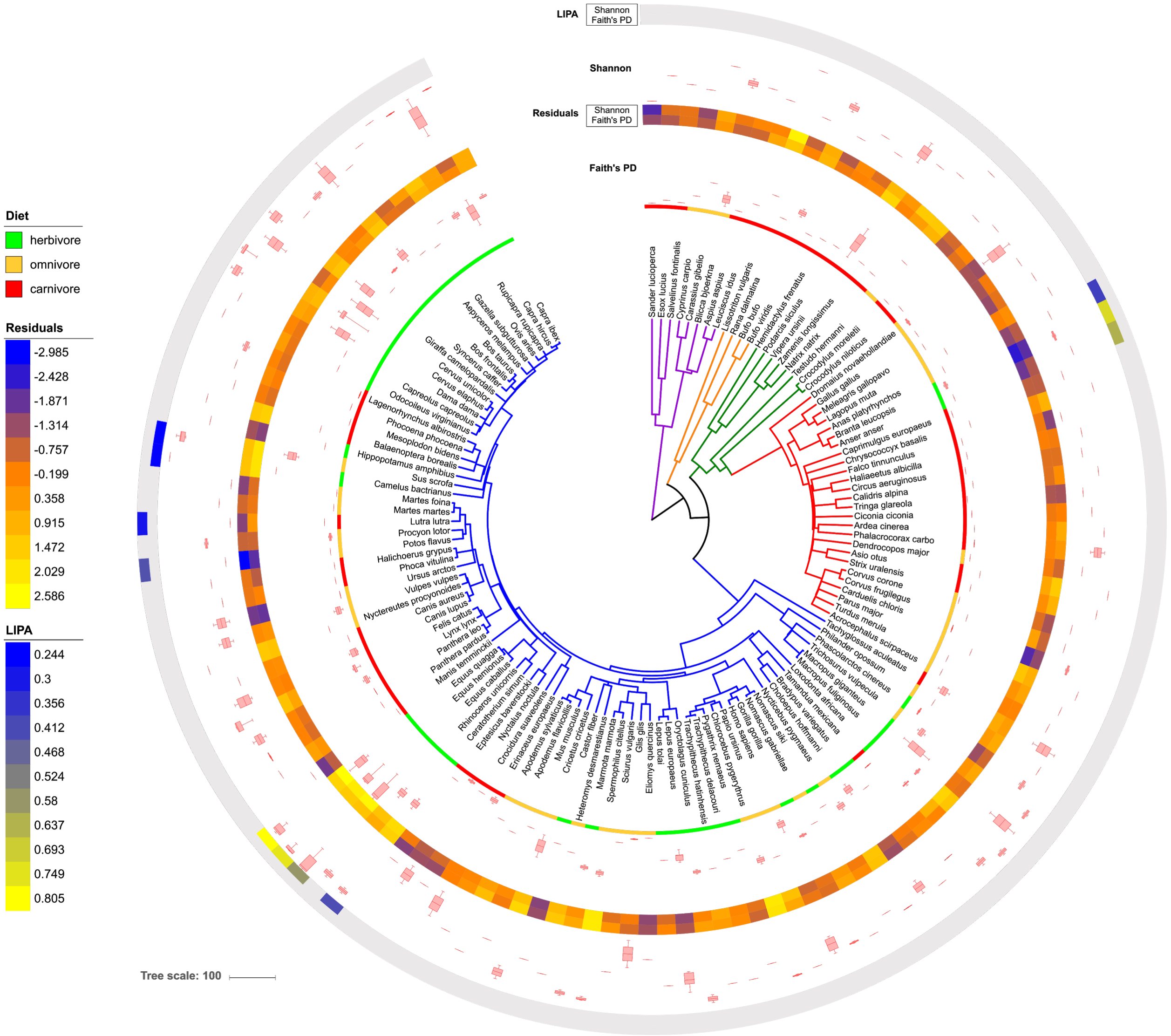


**Supplementary Fig. 10**. *Very little phylogenetic signal of alpha-diversity after accounting for diet.* The phylogeny is the same as shown in Fig. 1. The boxplots show distributions of alpha-diversity values (inner = Faith's PD; outer = Shannon index) among samples for the same host species. The heatmap between the boxplots shows residuals for both alpha-diversity measures after regressing out diet (all diet components). The outer heatmap shows LIPA Moran's I index values (ie., local phylogenetic signal), with grey representing all non-significant (significant defined as Adj. P-value <0.05).

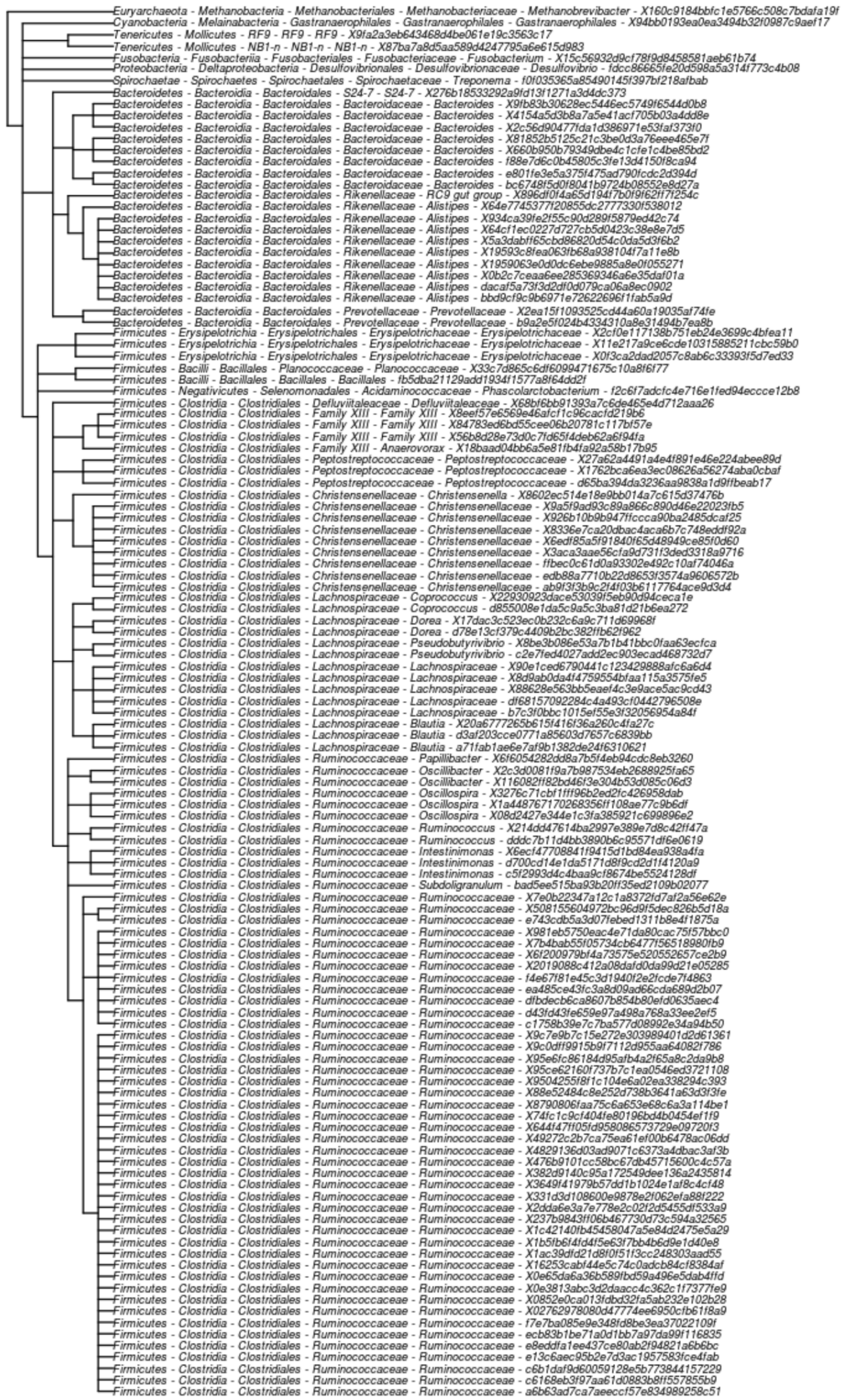


**Supplementary Fig. 11**. The cladogram is the same as in Fig. 4, but tip labels show the full taxonomy of each OTU: “phylum - class - order - family - genus - OTU-ID”.

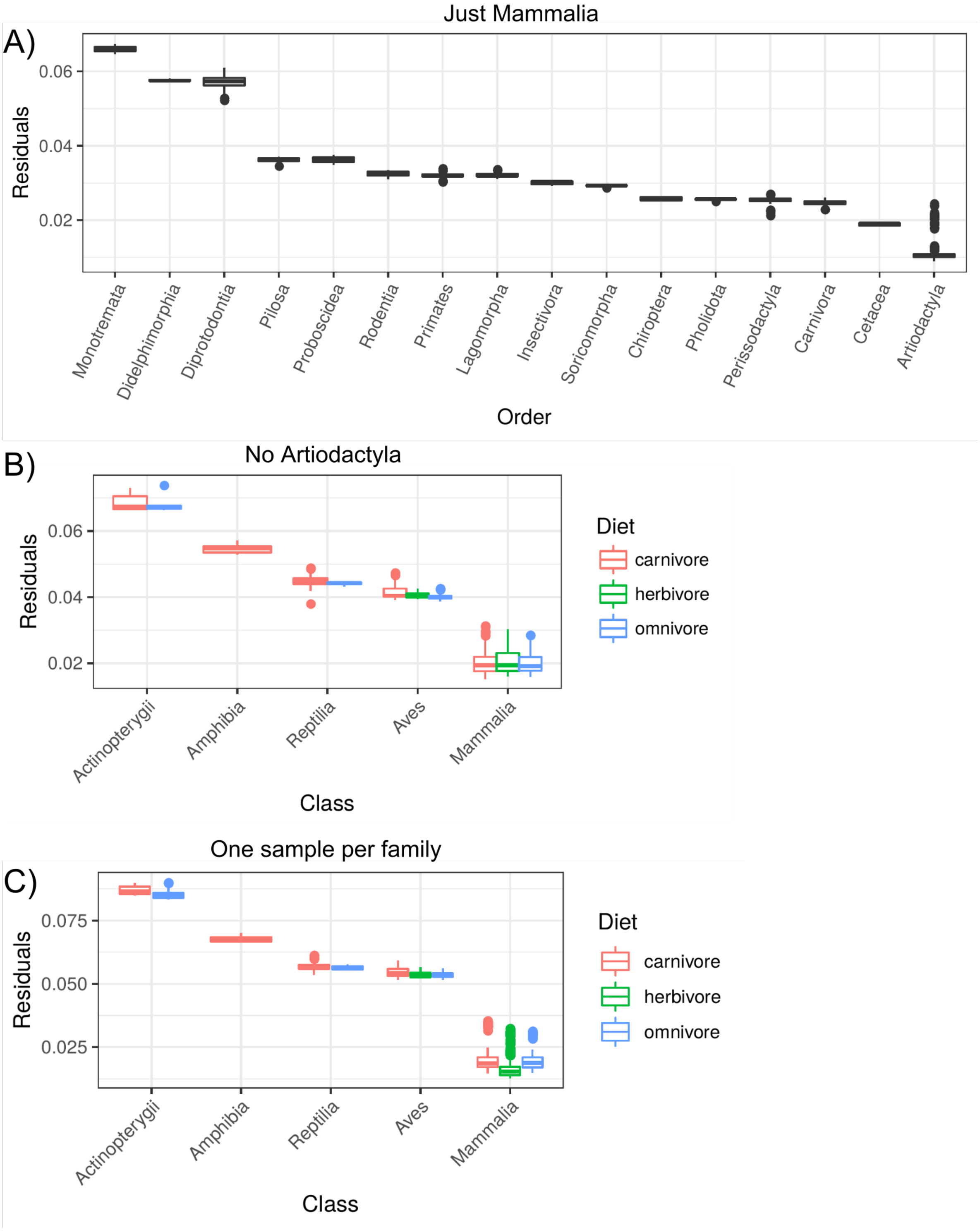


**Supplementary Fig. 12.** *The PACo test of co-phylogeny is robust to removing portions of the dataset*. The boxplots show Procrustean residuals for each host clade (smaller residuals means a better fit). PACo was performed on dataset subsets consisting of either A) just Mammalia hosts, B) Artiodactyla host removed, or C) just one sample per host family used.

A)

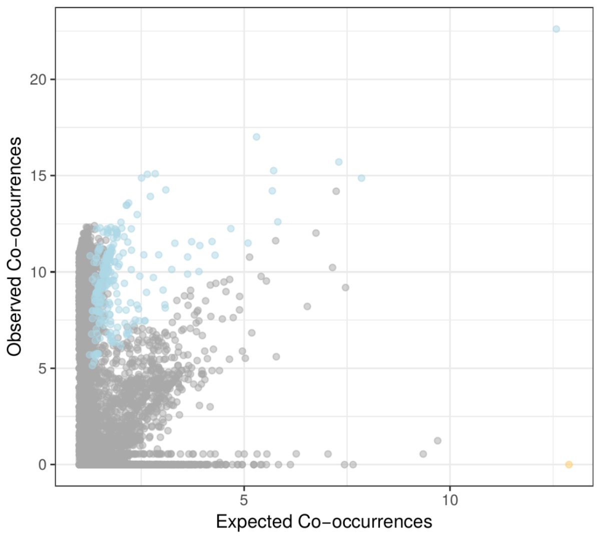


B)

| Phylum (x) | Phylum (y) | Sign | Perc. of Edges | Edges Norm. |
|---|---|---|---|---|
| Proteobacteria | Proteobacteria | -1 | 0.84 | 0.024 |
| Proteobacteria | Proteobacteria | 1 | 5.88 | 0.167 |
| Proteobacteria | Firmicutes | 1 | 1.28 | 0.112 |
| Firmicutes | Firmicutes | 1 | 1.07 | 0.276 |
| Firmicutes | Bacteroidetes | 1 | 1.06 | 0.014 |
| Bacteroidetes | Bacteroidetes | 1 | 0.90 | 0.037 |
| Proteobacteria | Bacteroidetes | 1 | 0.40 | 0.027 |
| Bacteroidetes | Firmicutes | 1 | 0.30 | 0.038 |
| Euryarchaeota | Firmicutes | 1 | 0.24 | 0.003 |

**Supplementary Fig. 13.** *Significant co-occurrences are mostly positive*. A) Most edges that significantly differed from null model expectations were positive co-occurrences. Each point signifies the expected versus observed co-occurrence among OTUs. B) The table lists the taxonomic composition of the significant edges, with “Sign” signifying a positive (1) or negative (-1) co-occurrence pattern, “Perc. of Edges” signifying the percent of total edges, and “Edges Norm.” signifying the fraction of edges normalized by the sum of nodes associated with those edges.

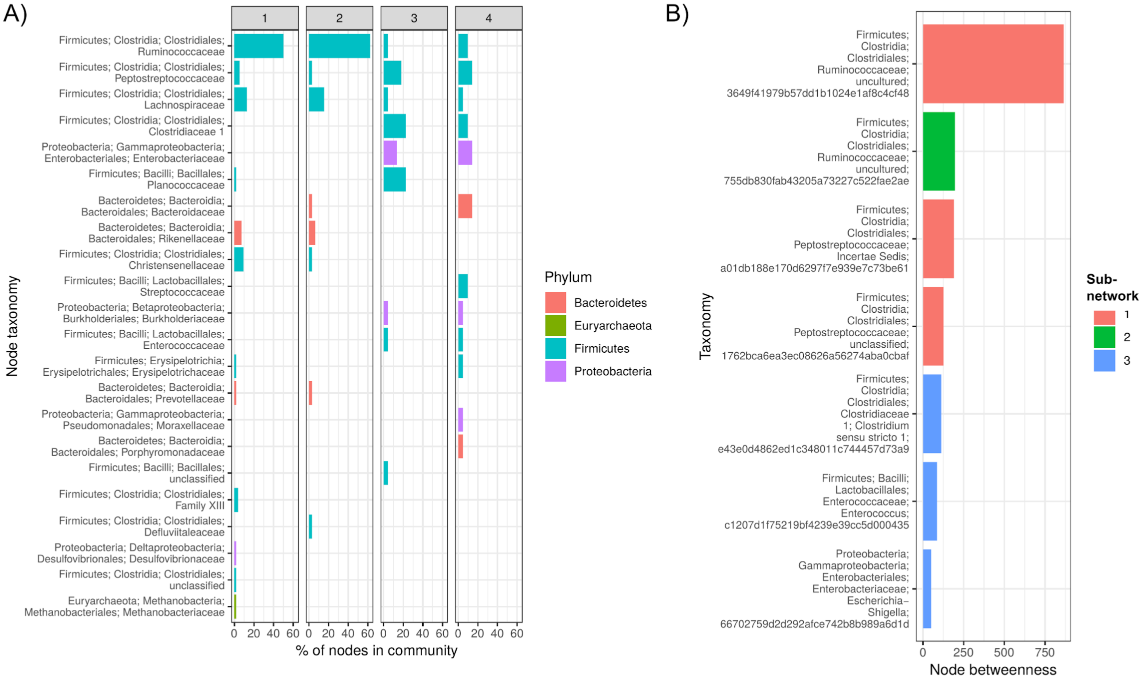


**Supplementary Fig. 14.** A) The bar chart shows the taxonomic composition of the nodes in each sub-network (x-axis plot facet) as defined in Fig. 7, with OTUs grouped at the genus level, and taxon labels listing “Phylum; Class; Order; Family; Genus”. B) The plot shows the centrality betweenness value for all OTUs with a value of >50, and taxon labels are coded as “Phylum; Class; Order; Family; Genus; OTU”.
